# Isolation of a nanobody specific to the PstS-1 protein and evaluation of its immunoreactivity with structural components of *Mycobacterium tuberculosis* granuloma

**DOI:** 10.1101/2025.06.05.658172

**Authors:** Yogesh P Dhekale, Kumarasamy Jothivel, Samar Kumar Ghorui, Gagan Deep Gupta, Shweta Singh, Nawab Singh Baghel, Savita Kulkarni, Pramod Kumar Gupta

## Abstract

*M. tuberculosis* (Mtb) causes infectious granulomatous disease tuberculosis (TB). Within TB granuloma, foci of Mtb secreted antigens anchored on the surface of either bacilli or host cells may serve as targetable biomarkers for antibody based molecular imaging of TB. Nanobody is better suited over conventional antibody or its fragment derivatives for molecular imaging due to its quick localization in target tissue and rapid clearance from off-target organs. Here, we report the production of a high affinity nanobody against PstS-1 protein of Mtb which helps bacilli in phosphate uptake as well as host cell adhesion. C8 nanobody was isolated from a phage display library displaying nanobodies which was constructed from a camel immunized with secretory proteins of Mtb. C8 Nb was characterized *in vitro* and *in vivo* for its immunoreactivity against PstS-1 protein. Ability of C8 Nb to bind PstS-1 protein present on the surface of Mtb bacilli or adhered on macrophages and its localization around BCG cells injected intramuscularly in mice demonstrate its potential in the development of molecular imaging for tuberculosis.

## Introduction

Tuberculosis (TB), caused by airborne pathogen *Mycobacterium tuberculosis* (Mtb), remains ancient and deadliest infectious disease of mankind. In the year 2023, WHO estimated 10.8 million new cases & 1.25 million deaths related to TB. Out of 10.8 million estimated cases, only 8.2 million TB patients were diagnosed; leaving behind a steadily narrowing global gap between the estimates and notifications of 2.6 million cases [1]. TB is a preventable as well as curable disease with proper medication, however, obstacles like variable efficacy of BCG vaccine (the only licensed vaccine), delay in decisive diagnosis, prolonged anti-TB drug therapy and emergence of drug resistance are affecting the rate of reduction in TB incidences and deaths, which obscures the goals of ‘End TB Strategy’ in high TB burden countries [2]. In the TB control programme, diagnosis is a key component as the early and precise diagnosis can promote prompt initiation of treatment to reduce further disease transmission, damage to infected tissue and development of drug resistance [3]. Algorithm of TB diagnosis prescribes various tandem tests depending on the outcome of tests and alignment of symptoms with tuberculosis. The cost-effective and rapid sputum microscopy is serving TB diagnosis for more than a century. However, the test suffers sensitivity when the bacillary load is less than 10^4^ bacilli/ml of sputum [4]. Chest X-ray examination is a very well-established primary diagnostic tool which provides anatomical information necessary for the management of TB suspects [5]. However, due to inconsistency in the extent of TB related tissue level abnormalities which is governed by competency of immune system [6] and their overlap with other clinical conditions [7], further examination is required to confirm the radiological findings suspecting TB. In the last decade, nucleic acid amplification-based diagnostic tests (NAATs) have revolutionized detection of Mtb and drug resistance with their greater specificity and sensitivity along with shorter turnaround time. However, smear-negative paucibacillary TB in immunocompromised, HIV co-infected and pediatric population [8, 9] is inadequately diagnosed by these tests, which is responsible for ∼16% of TB transmissions [10]. TB culture test, which has turnaround time of few weeks, remains the gold standard especially when rapid tests fail to detect Mtb. Despite plethora of available diagnostic tests, in year 2023, out of 8.2 million diagnosed TB patients, only 63% cases were bacteriologically confirmed, while 37% of TB patients were diagnosed clinically and initiated with empirical treatment to bring down the gap between TB case estimates and notifications to 2.6 million [1]. Therefore, new diagnostic modalities are urgently required to tackle the limitations of existing diagnostic methods.

Efforts of immune system to control Mtb growth and dissemination into distant organs produces hallmark lesion called granuloma, where wall formed by recruited neutrophils, macrophages, dendritic cells, monocytes, NK cells and lymphocytes surrounds free as well as phagocytosed Mtb bacilli [11]. The detection of TB granuloma has been explored using anatomical imaging through chest X-ray, ultrasound, MRI and CT, however, these techniques are not specific to TB [12, 13]. Active metabolism in Mtb bacilli as well as immune cells of granuloma has been explored as TB diagnostic biomarkers through minimally invasive PET/CT and SPECT imaging. Various radiolabelled tracers which are based on the precursor metabolites or antibiotic drugs have shown potential in detection of TB granuloma, however, their nonspecific nature is limiting their use only up to disease staging, treatment response monitoring and predicting relapse [14]. Structurally, granuloma is made up of aggregated immune cells surrounding Mtb bacilli at the core. Growing Mtb actively secretes various proteins which promote extracellular as well as intracellular growth of pathogen. These proteins accumulate in the granuloma through their interactions with pathogen cell wall or host cell surface receptors. Presuming sufficient concentration of accumulated antigens at granulomatous infection site, antibodies raised against such secreted antigens have been explored for molecular imaging of TB granuloma. In the earlier attempts of immunoscintigraphy, TB lesions were imaged using radiolabelled intact antibodies [15, 16] or F(ab’)_2_ fragments [17] specific to mycobacterial antigens. However, imaging contrast was produced only after days to weeks, post tracer administration. Molecular size of tracer determines its kinetics of accumulation in the target tissue and clearance from the background, which ultimately decides optimum timing for imaging [18]. Currently, various small sized recombinant proteins with faster pharmacokinetics are being developed for tumor imaging and therapy. A nanobody is ∼15kDa recombinant protein derived from variable domain of heavy chain only antibody produced by camelids. These camel derived nanobodies are less immunogenic due to their high sequence homology with human VH, which have advantage over monoclonal antibodies in terms of their repeated administrations in the patients. In contrast to intact IgG antibodies, which remain in the circulation for longer period of time due to their slow extravasation and binding with Fcγ receptor, compact nanobodies offers better tissue penetration and faster clearance from the circulation to produce imaging contrast within few hours of the administration [18, 19]. This feature of nanobody will help in diagnosing extrapulmonary TB including TB meningitis, as nanobodies have shown the ability to cross the blood brain barrier [20]. Also, use of recombinant antibodies lacking Fc domain mitigates their nonspecific accumulation around the pathogen or host cells due to the interactions between Fc domain and Fcγ receptors [21–23]. Use of pathogen specific antibodies may help in discriminating infection from sterile inflammations and malignancies.

Cell wall associated PstS-1 protein of Mtb is a phosphate binding subunit of Phosphate Specific Transport (PST) system [24–27]. In the Mtb cell wall, PstS-1 protein remains anchored through its hydrophobic tail and participates in the phosphate transport [28, 29]. Apart from this, PstS-1 protein is also known to play role in Mtb infection through its interactions with various pattern recognition receptors (PRR) including mannose receptor (MR), TLR2 and TLR4 [27, 30, 31]. Therefore, PstS-1 protein capable of accumulating in the TB granuloma through its interactions with Mtb cell wall or PRRs expressed on the immune cells forming granuloma appears to be a suitable target for nanobody based molecular imaging of TB granuloma.

In the present study, we have constructed a phage displayed nanobody library (henceforth mentioned as nanobody library) and used for isolation of nanobodies against PstS-1 protein of Mtb. Binding specificity, affinity and ability of lead C8 nanobody (C8 Nb) to bind PstS-1 protein present on the surface of Mtb bacilli or macrophages were evaluated using *in vitro* binding assays. Further confirmation of *in vivo* localization of ^125^I-C8 Nb around BCG cells injected through intramuscular route in mouse model shows that the C8 Nb may be useful in the development of molecular imaging for precise diagnosis and staging as well as evaluation of treatment response in tuberculosis.

## Results

### Construction of Anti-Mtb nanobody library

To construct anti-Mtb nanobody display library, a healthy dromedary was immunized with mixture of Mtb secretory proteins. Figure 1 depicts the schematic map for construction of anti-Mtb nanobody display library and isolation of anti-PstS-1 C8 Nb. Briefly, after the 4^th^ booster, camel seroconversion was confirmed by ELISA, where post immunization camel serum showed generation of heavy chain only antibodies specific to PstS-1 protein (Figure 2A). cDNA prepared using total RNA extracted from camel PBMCs was used for nanobody gene preparation in two step PCR. IgG-PCR produced mixture of amplicons of gene fragments (900bp, 690bp and 620bp) was electrophoresed on agarose gel and 690bp as well as 620bp amplicons were purified (Figure 2B). These purified amplicons were used as DNA template in V_H_H PCR, which produced smeared band of ∼400bp-500bp size fragments (Figure 2C). EcoRI & HindIII modified nanobody gene amplicons were ligated to T7 phage vector DNA arms and insert-vector ligation was confirmed by PCR (Figure 2D). Subsequent *in vitro* phage packaging reaction yielded a library of ∼2.5×10^5^ nanobody clones (Figure 2E).

**Figure 1.**
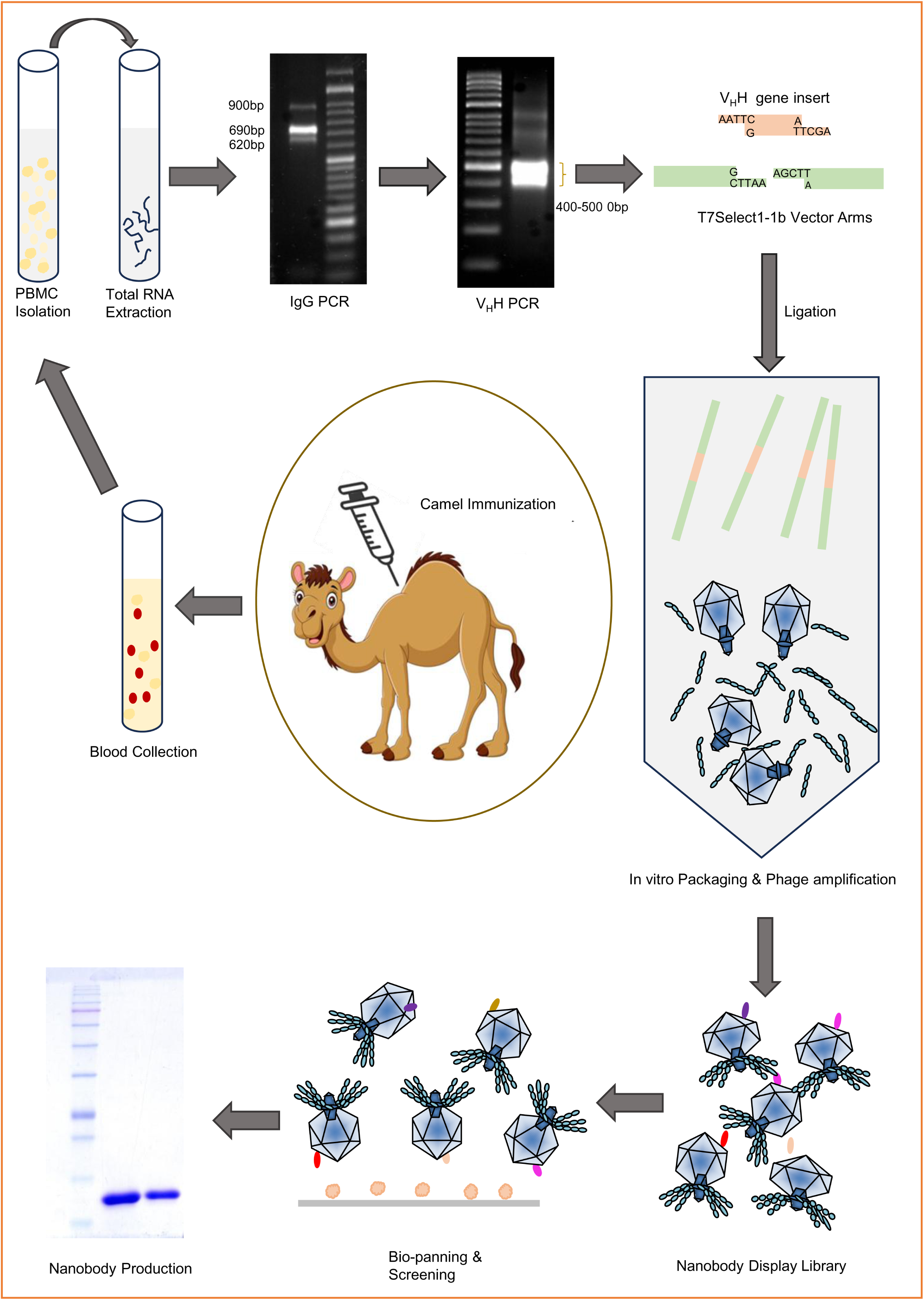
Schematic map for construction of anti-Mtb nanobody display library and isolation of anti-PstS-1 C8 Nb.

**Figure 2.**
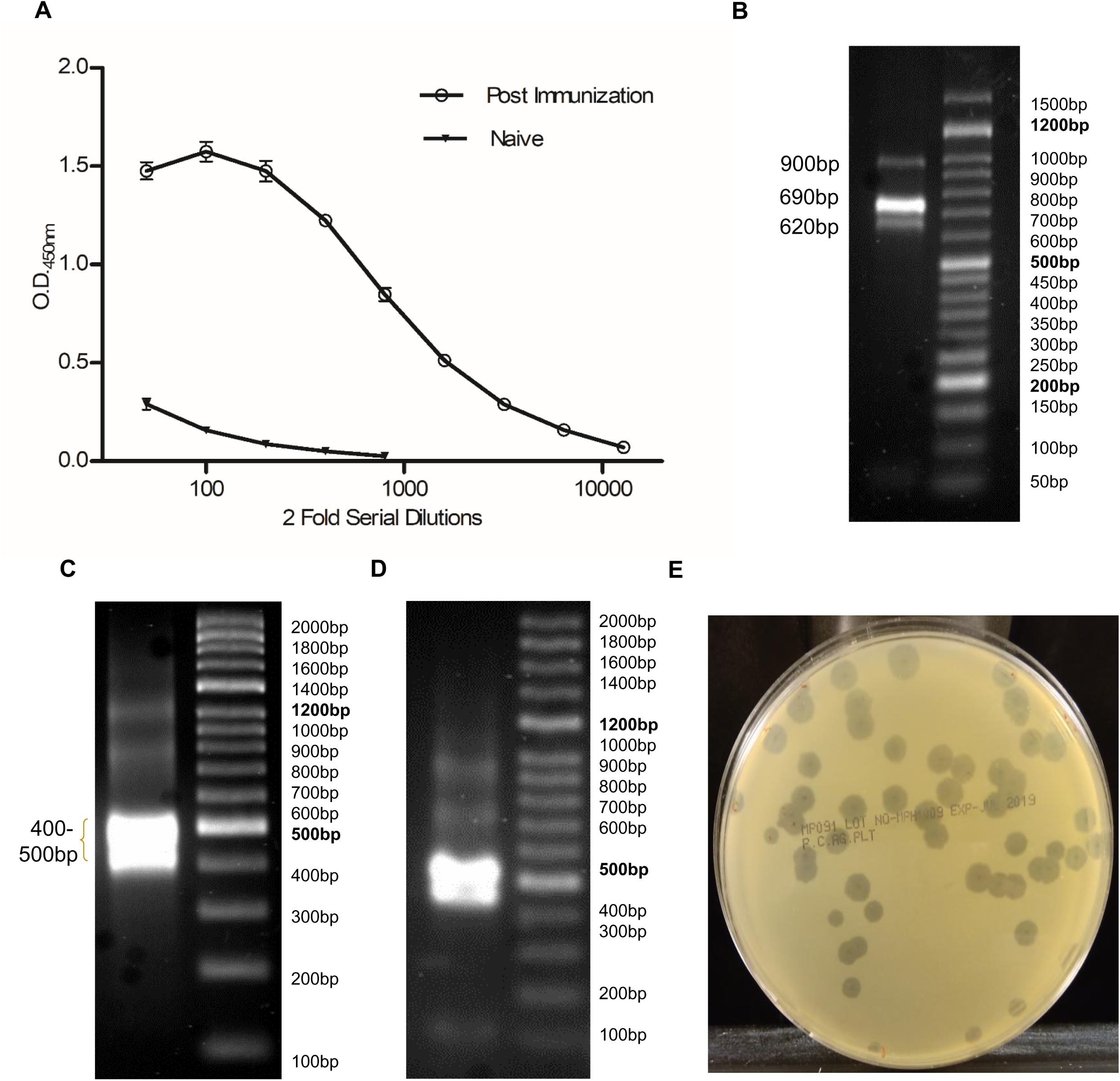
Nanobody display library construction. A) Camel seroconversion was confirmed by ELISA, where ELISA wells coated with rPstS-1 protein were treated with serially diluted camel serum and bound antibodies were measured by incubating rabbit anti-camelid V_H_H antibody-HRP conjugate. B) In IgG PCR, genetic locus between FR1 region of variable domain and 5’ end of CH2 domain were amplified, which yielded 620bp, 690bp and 900bp size amplicons. corresponding to IgG3, IgG2 and IgG1 subclasses, respectively. C) In V_H_H PCR, primers amplified the region between FR1 and FR4, which produced smeared band of ∼400bp-500bp size amplicons. D) Ligation of nanobody gene insert and T7 vector DNA arms was confirmed by PCR using vector based primers (T7SelectUp and T7SelectDown). E) Nanobody library size (2.5×10^5^ clones) was measured by PFU assay.

### Isolation of PstS-1 protein specific nanobodies

Primary nanobody library was amplified in liquid culture and used to isolate nanobodies specific to rPstS-1 protein which was overexpressed and purified from *E. coli* BL21(DE3) in soluble form (Figure 3A). In bio-panning, nanobody library was enriched >100 fold against coated rPstS-1 protein (Figure 3B). After enrichment, immunoreactivity of randomly selected 30 phage clones was assessed in two steps. In the first assay, positive binding was observed for 29 nanobody clones against coated rPstS-1 protein (Figure 3C). In the subsequent assay, which was performed to ensure that the binding obtained in first assay was specific to rPstS-1 protein rather than other well components, 10 selected phage lysates captured through phage tail reproduced immunoreactivity with ^125^I-labelled rPstS-1 protein (Figure 3D). Further, sanger sequencing revealed that nanobody clone 8 and 11 were identical at nucleotide sequence, while clone 5 had two transversions in FR1 region as compared to clone 8/11(C8 Nb). C8 Nb was selected for the further studies and its version carrying C-terminal 6xHis tag was overexpressed in *E. coli* BL21(DE3) in soluble form. Subsequently, nanobody was purified to the single band using metal affinity chromatography, anion exchange chromatography and gel filtration sequentially (Figure 3E).

**Figure 3.**
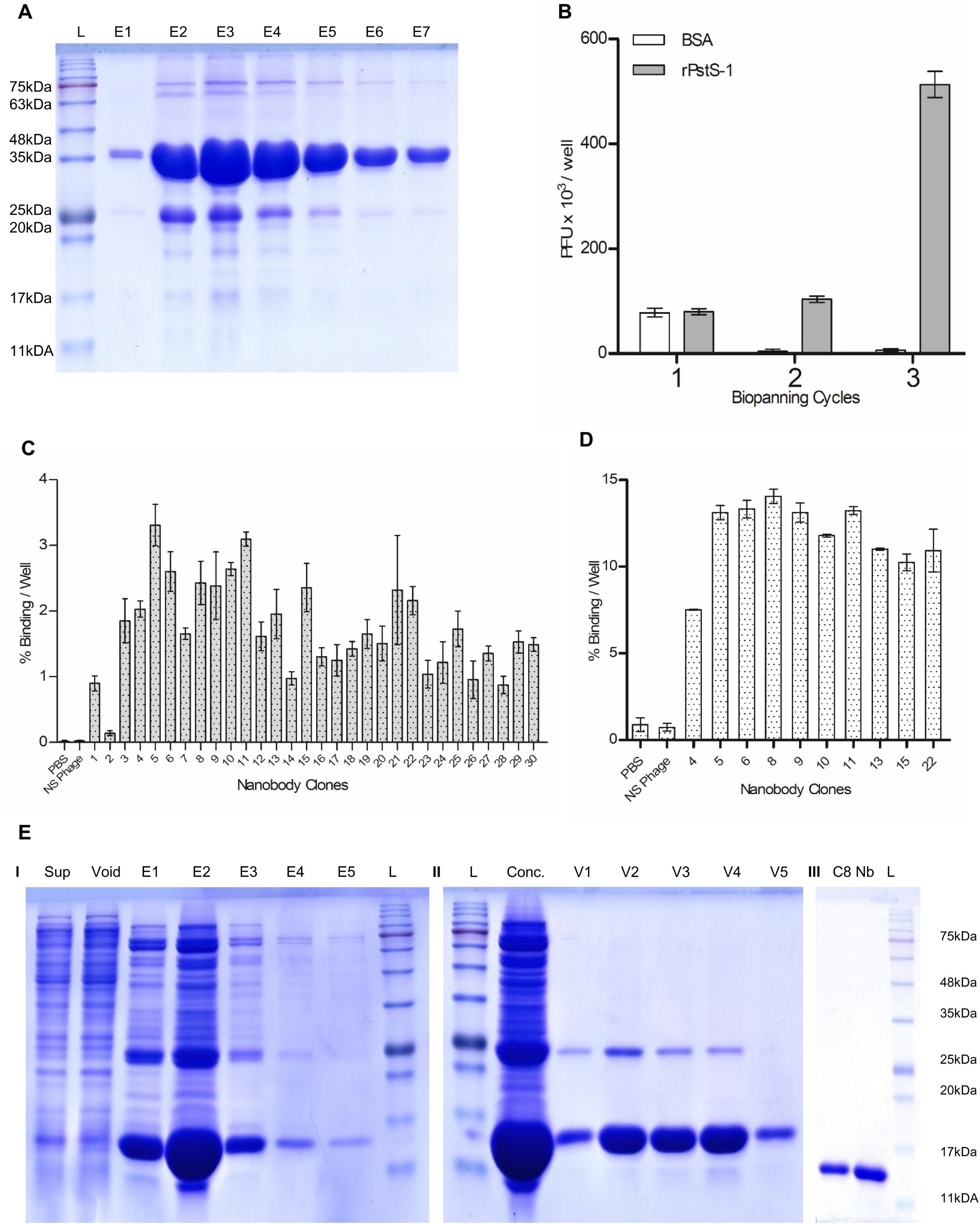
Isolation of nanobody against PstS-1 protein of Mtb. A) rPstS-1 protein of Mtb was overexpressed in *E. coli* BL21(DE3) and soluble protein was purified using Ni-NTA agarose column chromatography. B) Three cycle biopanning was performed for enrichment of PstS-1 protein specific nanobodies. Antigen specific nanobody enrichment was confirmed by PFU assay. C) Nanobody screening was carried out using two assay formats. Immunoreactivity of randomly selected nanobody clones was evaluated by reacting monoclonal phage lysates with coated rPstS-1 protein and bound phage were measured by incubating ^125^I-labelled anti-T7TF mAb. D) Reproducibility of immunoreactivity of nanobody clones which were selected in the first assay was confirmed by capturing phages in anti-T7TF mAb coated wells, followed by reacting wells with ^125^I labelled rPstS-1 protein. Nanobody clones giving binding in both assays were selected for DNA sequencing. E) C8 Nb was overexpressed in *E. coli* BL21(DE3) and soluble protein was purified using I) Ni-NTA agarose column chromatography and eluates containing nanobody were pooled together and concentrated and subjected to II) DEAE cellulose column chromatography. III) Void fractions containing unbound nanobody were pooled together and purified on Superdex^TM^ G75 Increase 10/300 GL column.

### High affinity C8 Nb specifically recognizes PstS-1 protein

Nanobody clone enrichment and screening were performed on rPstS-1 protein, which carries vector derived additional amino acid residues including 6xHis tag. Also, being secretory in nature, PstS-1 protein may share consensus amino acid sequences involved in protein trafficking and interactions with cell wall components [32]. Therefore, to ascertain that C8 Nb is specifically recognizing PstS-1 protein but no other proteins from Mtb lysate; we probed rPsts-1 protein as well as crude protein preparations of Mtb, BCG, *M. smegmatis* and *E. coli* blotted on PVDF membrane with nanobody (Figure 4A). After sequentially incubating the membrane with C8 Nb, rabbit anti-camelid V_H_H antibody-HRP conjugate and chemiluminescent substrate, an intense band of ∼38kDa size was developed in the lane of rPstS-1 protein. Similarly, single band was visible around similar size in the lanes loaded with crude protein preparations of Mtb and BCG. In the lanes representing protein preparations from either *E. coli* or *M. smegmatis* there was no development of corresponding size band. In the lane of rPstS-1 protein, another faint band appeared near 75kDa size marker, which was missing in other lanes, could be a dimmer of rPstS-1 protein. Collectively immunoblotting data confirm that C8 Nb specifically recognizes PstS-1 protein from crude protein preparations of Mtb and BCG.

**Figure 4.**
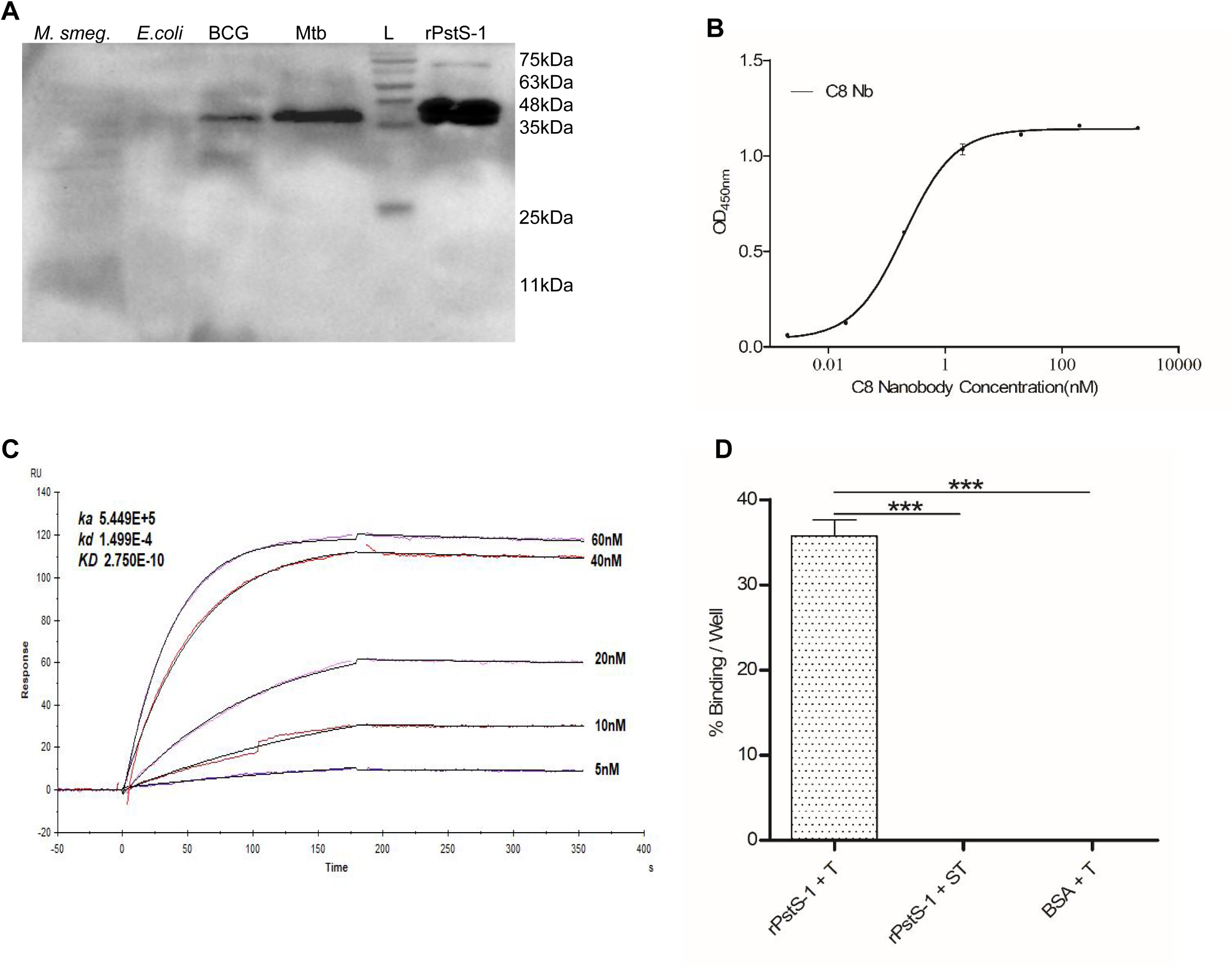
High affinity C8 Nb specifically recognizes PstS-1 protein. A) Crude mixture of proteins or rPstS-1 protein were electrophoresed on SDS-PAGE and blotted on PVDF membrane, followed by sequential incubation with C8 Nb and rabbit anti-camelid V_H_H-HRP conjugate. C8 Nb recognized PstS-1 protein of Mtb and homologue expressed by *M. bovis* BCG. B) Saturation binding ELISA showed 0.198nM equilibrium dissociation constant of C8 Nb. C) Ka (5.44×10^5^ M^−1^ sec^−1^), Kd (1.499×10^-4^ sec^−1^) and K_D_ (0.275×10^-9^M) values of C8 Nb were measured using Biacore X100 SPR system (GE Healthcare). D) Immunoreactivity of ^125^I-C8 Nb was confirmed against rPstS-1 protein coated in ELISA wells by radioimmunoassay. All values are mean ± s.d. Significance was calculated by one-way ANOVA followed by Dunnett’s multiple comparisons post hoc test; ***P <0.001. Data are representative of more than three independent experiments.

Subsequently, affinity of C8 Nb was evaluated by measuring equilibrium dissociation constant (K_D_) by saturation binding ELISA as well as SPR. In ELISA, rPstS-1 protein coated wells were sequentially incubated with serially diluted C8 Nb, rabbit anti-camelid V_H_H antibody-HRP conjugate and TMB substrate. ELISA results demonstrated that the K_D_ value is 0.198nM (Figure 4B). Further, C8 Nb and PstS-1 antigen binding kinetics were also characterized using SPR on Biacore X100 SPR system (GE Healthcare), where 1:1 binding model revealed that C8 Nb has Ka, Kd and K_D_ values of 5.44×10^5^M^−1^sec^−1^, 1.499×10^-4^sec^−1^ and 0.275nM, respectively (Figure 4C). Sub-nanomolar K_D_ value obtained in the both binding experiments confirmed that C8 Nb has sufficient affinity for the purpose of molecular imaging.

Further, radioimmunoassay (RIA) was performed to confirm the immunoreactivity of ^125^I labelled C8 Nb against rPstS-1 protein (Figure 4D). We chose ^125^I radionuclide for nanobody labelling due to the ease of labelling procedure and absence of particulate emissions. C8 Nb was labelled with ^125^I radionuclide using Iodogen method and purified using size exclusion chromatography. When ^125^I-C8 Nb (tracer) was incubated in wells coated with rPstS-1 protein, there was 35% tracer binding whereas in BSA coated control wells, tracer binding was <1%. Moreover, when the antigen coated wells were reacted with tracer which was spiked with 1000-fold excess unlabelled C8 Nb, tracer binding was reduced to <1% of added tracer, which confirms that immunoreactivity of C8 Nb was retained after radioiodination.

### C8 Nb recognizes PstS-1 protein present on the surface of drug sensitive as well as MDR strains of Mtb

PstS-1 protein in TB granuloma, adhered on the surface of either Mtb bacilli or macrophages, may serve as antigenic target for C8 Nb. In order to investigate whether epitope on PstS-1 protein which is adhered on either Mtb bacilli or macrophages is accessible for C8 Nb binding we analysed immunoreactivity of ^125^I-C8 Nb with Mtb bacilli; as well as macrophages treated with antigen. When the tracer was incubated with bacilli prepared from log phase cultures, pellet of Mtb and BCG retained 20.82% ± 0.56 and 14.62% ± 2.36 of added tracer respectively, while there was <0.5% tracer retention in the pellets of *M. smegmatis* and *E. coli*. Additionally, when the similar bacterial pellets were treated with tracer mixed with 1000-fold excess unlabelled C8 Nb, the tracer binding was reduced to <1% of added quantity (Figure 5A). Binding assay results confirm that epitope of PstS-1 protein anchored on the surface of Mtb and BCG bacilli is accessible for C8 Nb binding, moreover, this nanobody did not recognise PstS-1 protein homologues expressed by *M. smegmatis* as well as *E. coli.* In the similar experiment, immunoreactivity of ^125^I-C8 Nb with two multi drug-resistant clinical isolates of Mtb belonging to different genetic lineages such as LAM (Latin American, Mtb LAM) and Beijing (Mtb Beijing) was confirmed and data suggested the binding of C8 Nb on the surface of both the Mtb isolates (Figure 5B).

**Figure 5.**
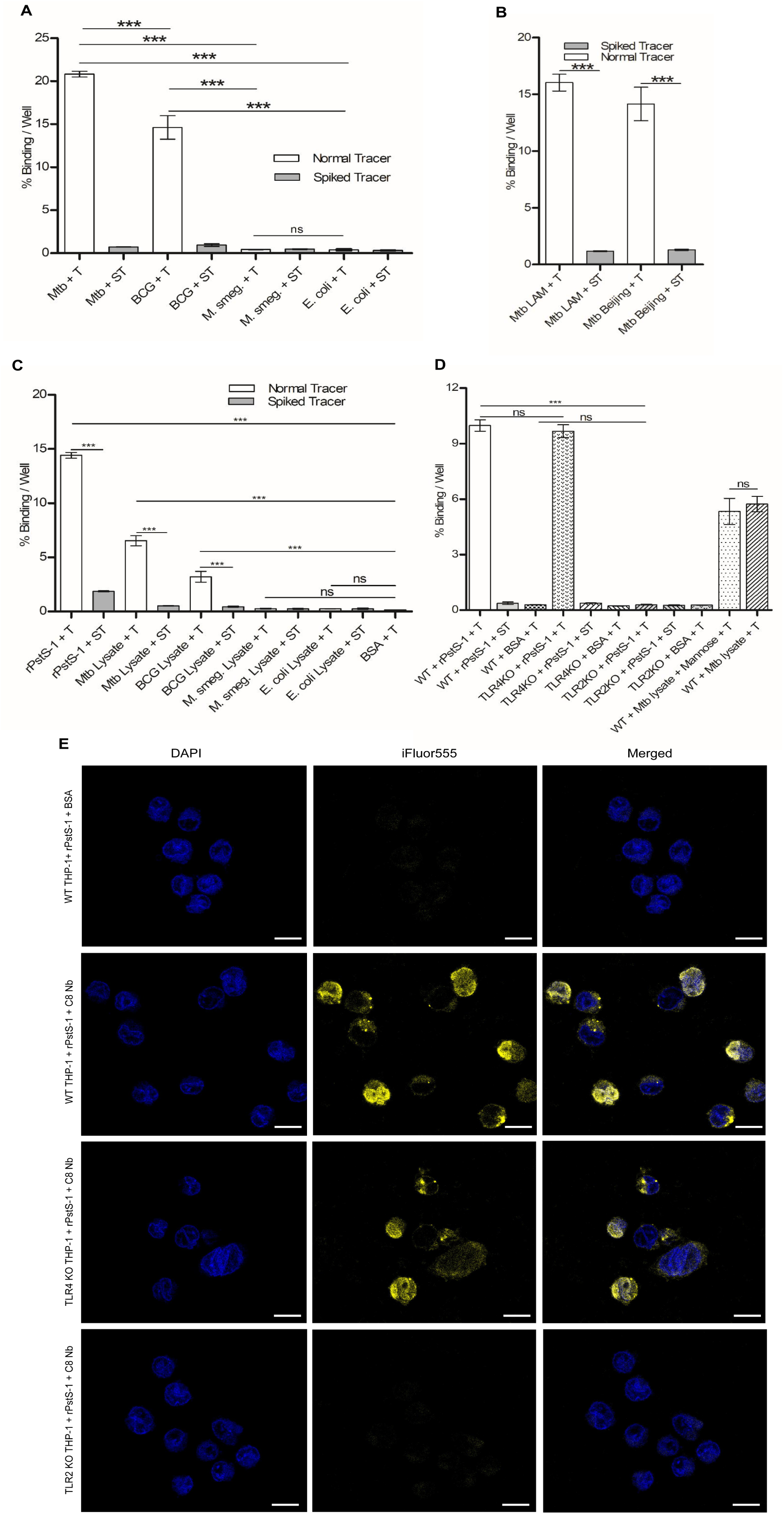
Immunoreactivity of C8 Nb with PstS-1 protein adhered on the surface of either mycobacteria or macrophages. A) When single cell suspensions of various bacteria were probed with ^125^I-C8 Nb, only Mtb and BCG pellets showed tracer binding. B) Immunoreactivity of ^125^I-C8 Nb was confirmed with bacilli of Mtb LAM and Mtb Beijing strains. C) Protein mixtures prepared from various bacterial lysates were incubated with formaldehyde fixed RAW 264.7 cells and then cells were incubated with ^125^I-C8 Nb. Tracer showed binding with the cells which were treated with either rPstS-1 protein or protein lysates prepared from Mtb or BCG bacilli. In all assays, specific binding of ^125^I-C8 Nb was confirmed by including spiked tracer which was prepared by mixing the tracer with excess unlabelled C8 Nb. D) Binding studies with THP-1 macrophages revealed that the epitope on PstS-1 protein bound to only TLR2 is accessible to C8 Nb for binding, while MR and TLR4 are not playing role in binding. E) Fluorescence microscopy images showing TLR2 involved in C8 Nb binding with PstS-1 protein treated THP-1 macrophages. Scale bars: 10μm. For panel A-D, all values are mean ± s.d. derived from more than three independent experiments. Significance was calculated by one-way ANOVA followed by Dunnett’s multiple comparisons or Bonferroni’s post hoc test; ***P <0.001.

### C8 Nb recognises PstS-1 protein adhered on the surface of macrophages

Along with Mtb bacilli, immune cells surrounding the Mtb foci at infection site are major structural components of TB granuloma. Immune cells use various PRR to recognise pathogen associated molecular patterns. PstS-1 protein of Mtb is known to bind PRRs such as TLR2, TLR4 and MR which are expressed on the surface of immune cells. This assay was performed to assess whether the PstS-1 protein bound to PRR on RAW 264.7 macrophages could be recognized by C8 Nb. When formaldehyde fixed macrophages were treated with either rPstS-1 protein or bacterial crude protein mixtures and ^125^I-C8 Nb, cells treated with rPstS-1 protein, Mtb protein mixture and BCG protein mixture retained 14.42% ± 0.36, 6.54% ± 0.67 and 3.2% ± 0.71 of added tracer, respectively, which confirms that PstS-1 protein bound to macrophage surface receptors can be recognized by C8 Nb (Figure 5C). Control wells, wherein cells were incubated with protein preparation from *E. coli, M. smegmatis* or BSA retained <0.5% of added tracer. Interestingly, in the wells where cells were treated with similar protein preparations and the tracer which was spiked with 1000-fold excess unlabelled C8 Nb, the tracer binding was reduced to <0.5%. Therefore, our data confirmed the specific binding of C8 Nb to the PstS-1 protein anchored on the macrophages.

### C8 Nb recognizes PstS-1 protein bound to TLR2 but not to MR and TLR4

To understand the PRR involved in binding with macrophages, we used wild type (WT) as well as TLR2/4 knockout (KO) human monocytic cell line THP-1 (Figure 5D). PMA activated THP-1 cells were fixed with formaldehyde, followed by sequential incubation with rPstS-1 protein and ^125^I-C8 Nb. WT THP-1 cells expressing all the three PRR retained 9.98% ± 0.44 of added tracer, which was comparable with 9.68% ± 0.50 tracer binding with TLR4 KO cells. Similarly, WT THP-1 cells treated with mixture of Mtb crude proteins showed 5.34% ± 1.01 and 5.73% ± 0.58 of tracer binding without any significant effect of presence or absence of added mannose in the protein preparations, respectively. Additionally, when similar antigen treated cells were incubated with tracer spiked with unlabelled C8 Nb, the tracer binding was reduced to the level of binding obtained with BSA treatment. These results confirmed that both MR as well as TLP4 are not involved in the binding. However, in the case of TLR2 KO cells, tracer binding was reduced to the level of binding obtained with BSA treatment, suggesting the involvement of TLR2 in binding with PstS-1 protein-C8 Nb complex. Role of TLR2 receptor in the binding of PstS-1 protein-C8 Nb complex on macrophages was also confirmed by confocal microscopy (Figure 5E). Wherein, surface of wild type and TLR4 KO THP-1 cells treated with rPstS-1 protein and C8 Nb showed binding of rabbit anti-camelid V_H_H cocktail-iFluor555 to a similar extent, while similarly treated TLR2 KO THP-1 cells showed minimal binding of rabbit anti-camelid V_H_H cocktail-iFluor555 which was comparable with WT THP-1 cells which were not treated with C8 Nb.

### ^125^I-C8 Nb localizes in intramuscularly injected *M. bovis* BCG cells in mouse

PstS-1 protein binding properties of ^125^I-C8 Nb were evaluated further in BALB/c mice which were injected with single cell suspensions of *M. bovis* BCG and *M. smegmatis* through intramuscular route (Figure 6A). In localization study (Figure 6B), right forelimb of mice injected with BCG retained 8427 ± 893 CPM/gm of tissue, which was significantly higher than the 2917 ± 503 CPM/gm of tissue retained in the left forelimb carrying *M. smegmatis*. Radioactivity counts obtained in the hindlimbs were lowest than any forelimb carrying bacterial pellets and were used for correcting the non-specific background counts.

**Figure 6.**
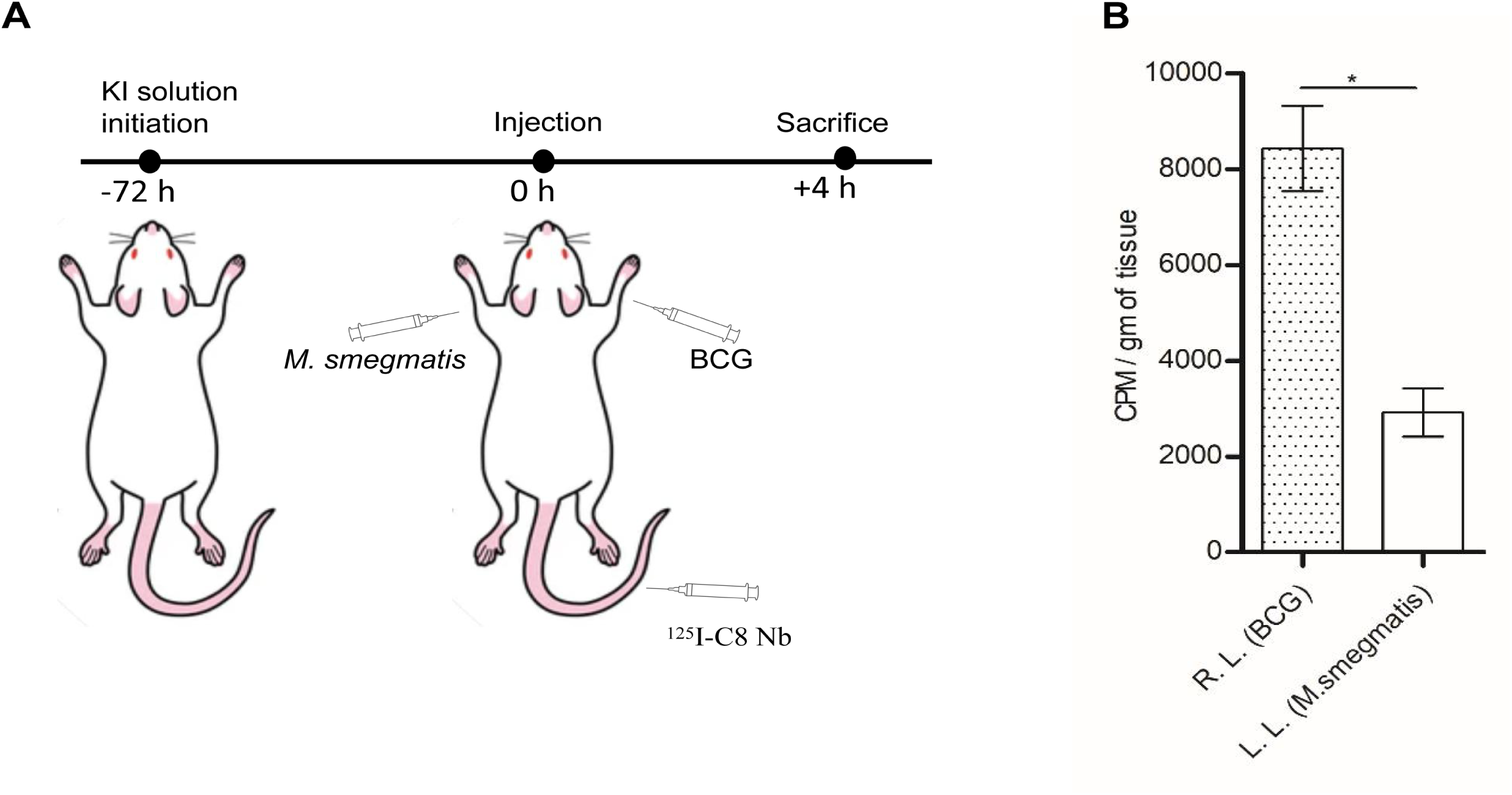
In vivo localization of ^125^I-C8 Nb in mycobacterial cells. A) To reduce uptake of free ^125^I in thyroid gland, BALB/c mice were fed with KI solution. For in vivo localization study, animals (n = 3 animals) were injected with single cell suspensions of BCG (right forelimb) and *M. smegmatis* (left forelimb) and then 25.91µCi of ^125^I-C8 Nb was administered through catheterized tail vein. B) Four hours post tracer administration, animal limbs were dissected and accumulated activity was measured in γ-photon counter. All mean ± s.d. values are derived from more than three independent experiments. Significance was calculated by one tailed T-test; *P <0.05.

## Discussion

Early diagnosis is one of the key components to reduce global epidemic of tuberculosis to an endemic disease under ‘End TB strategy’[2]. TB diagnosis includes multiple *in vitro* tests such as smear microscopy, microbial culture and various nucleic acid amplification based tests (NAATs) which offers inadequate sensitivity in samples with low bacillary load associated with paucibacillary and extrapulmonary TB [33, 34]. Along with the tests which detect Mtb bacilli, other tests such as tuberculin skin test and interferon-gamma release assays (IGRA), which measure memory T cell response generated during the Mtb infection are less sensitive in immunocompromised individuals [35, 36]. Quite the opposite, presence of humoral immune response against mycobacterial lipoarabinomannan is known to reduce the sensitivity of lateral flow immunochromatography in HIV negative TB patients [37]. In parallel to *in vitro* tests, non-invasive anatomical as well as molecular imaging modalities have been pursued for TB diagnosis. In the TB granuloma, foci of accumulated antigens secreted by Mtb poses itself as suitable target for antibody based molecular imaging of TB. Around the last decade of 20^th^ century, radiolabelled antibodies specific to Mtb antigens were evaluated for immunoscintigraphic detection of TB granuloma. Hazra *et al*. carried out immunoscintigraphical imaging of subcutaneously established Mtb infection in mouse using radiolabelled anti-PstS-1 protein monoclonal antibody [16]. In the similar study, Malpani *et al.* observed the earliest localization of ^131^I labelled anti-*M. bovis* BCG polyclonal antibodies in subcutaneous Mtb nodule after 3 days of intravenous injection, however, the best imaging contrast was developed only after 6 days [15]. When, Lee *et al.* injected ^131^I labelled polyclonal anti-*M. bovis* BCG F(ab)_2_ fragments, imaging of tuberculous lesion near knee joint was possible within 24 h post tracer administration due to faster clearance of tracer from the circulation [17]. These initial studies showed the possibility of radiolabelled antibody based imaging of extrapulmonary tuberculosis, however the specific and rapid Mtb detection using recently invented PCR technique waned the further interest in the exploration of molecular imaging for management of TB patients. Rejuvenated interest, after realizing the urgent need of diagnostic tool for the management of paucibacillary and extrapulmonary TB, has led to the development and evaluation of radiotracers like ^18^F-fluoro-deoxyglucose (FDG) [38], ^18^F-fluoro-deoxytrehalose (FTD) [39], and ^99^Tc-Ethambutol [40] for TB diagnosis. However, nonspecific nature of these molecules warrants further evaluation of Mtb antigen specific small antibody fragments which are known to quickly localize in target tissue and clear from the circulation which allows imaging within 2-4 h.

In the present study, we report the isolation of a PstS-1 protein specific high affinity nanobody from anti-Mtb nanobody library and evaluation of its immunoreactivity with PstS-1 protein adhered on the surface of Mtb bacilli as well as macrophages which are the structural components of TB granuloma. With the goal of isolating and characterizing nanobodies specific to various antigenic targets having diagnostic potential in TB, a nanobody library against whole mixture of secreted proteins of Mtb was constructed. Criteria for selecting target antigen for granuloma imaging was that; the actively secreted protein of Mtb should remain adhered on the bacilli surface and if it dissociates from Mtb, then it should remain anchored on the surface receptors of host immune cells which forms granuloma. PstS-1 protein of Mtb is part of PST complex composed of various protein subunits such as integral inner membrane proteins (PstA and PstC), cytoplasmic ATP hydrolyzing subunit (PstB) and periplasmic phosphate specific transport phosphate binding proteins (PstS) [24, 25]. Mtb actively traffics PstS-1 protein (also known as 38-kDa antigen, PhoS, pab, Ag78, Ag5 or PhoS1), an immunodominant glyco-lipoprotein [26, 27], across the cell membrane where the protein remains bound to the cell wall [28] through its hydrophobic tail and participates in the phosphate transport [29]. As an adhesion factor, PstS-1 protein interacts with mannose receptor (MR) on the macrophages to promote phagocytosis of pathogen [27]. Interaction of PstS-1 protein with both TLR2 and TLR4 induces the ERK1/2 and p38 MAPK signalling in monocytes, which plays crucial role in TNF-α and IL-6 expression during mycobacterial infection [30, 31]. Considering its anchoring properties and 38kDa size which can accommodate simultaneous binding of anchoring receptor and nanobody, we selected PstS-1 protein as a potential target for imaging of TB granuloma and isolated a high affinity C8 Nb. PstS-1 protein is trafficked to periplasm using signal peptide, where it anchors itself in the cell wall through its hydrophobic tail, is also present in the other cell wall proteins of Mtb [31, 32]. By probing whole cell lysate of Mtb on immunoblot we confirmed that C8 Nb is recognizing specifically PstS-1 protein from the Mtb lysate. As the phosphorous is a very crucial macronutrient for growth, the PST complex appeared very early in the evolution and homologous systems are also reported in the other bacteria [41]. On the same immunoblot, along with PstS-1 protein of Mtb, C8 Nb also recognized PstS-1 protein homologue expressed by *M. bovis* BCG, however, despite considerable similarity in amino acid sequence, there was no nanobody binding and band development around the corresponding size in the lanes of *E. coli* and *M. smegmatis* [32, 42], suggesting the absence of epitope recognized by C8 Nb. As the T7 phage library was grown on *E. coli* Rosetta-gami B5615 strain, nanobodies recognising antigens expressed by host cell may have got neutralised while phage growth. To assess the immunoreactivity of C8 Nb with PstS-1 protein homologues expressed by other clinically important mycobacteria including other members of MTB complex and non tuberculous mycobacteria was beyond the scope of this work, however, it could be done indirectly by epitope mapping through peptide microarray or Cryo-TEM.

C8 Nb has 6 tyrosine residues which are dispersed exclusively in the framework region-3. We labelled the purified C8 Nb with ^125^I using Iodogen method which attaches iodine atoms on the meta-position of benzene ring in tyrosine residues. When radiolabelled C8 Nb was evaluated for its binding with bacterial single cell suspensions, pellets of Mtb and BCG retained 20.82% ± 0.56 and 14.62% ± 2.36 of added tracer, respectively. This binding confirmed that the epitope on PstS-1 protein which is present in mycobacterial cell wall is accessible for nanobody binding and hence imaging of Mtb bacilli or clumps in the granuloma is possible. Less tracer retention in the pellets of *E. coli* and *M. smegmatis* shows less or no binding of tracer which is aligning with the results of immunoblot. Incubation of cell pellets with tracer spiked with excess unlabelled C8 Nb reduced the binding significantly, which confirms that the tracer binding with bacilli is antigen specific. This nanobody also detected PstS-1 protein which is present on the surface of multi drug-resistant clinical isolates such as Mtb LAM and Mtb Beijing, which shows wide range diagnostic usability of C8 Nb.

The immune cells forming wall of granuloma, which keeps Mtb from disseminating to other organs, express multiple PRRs including MR, TLR2 and TLR4 which are known to interact with PstS-1 protein [31]. In addition to detection of PstS-1 protein which is associated with Mtb cell wall, nanobody binding to antigen which is anchored on the surface of immune cells, will help in granuloma imaging by enhancing signal intensity. When RAW macrophages adhered with antigens were incubated with ^125^I-C8 Nb, cells from the wells treated with rPstS-1 protein retained 14.42% ± 0.36 of added tracer, while cells treated with protein mixture isolated from Mtb or BCG retained 6.54% ± 0.67 and 3.2% ± 0.71 of added tracer, respectively, which confirms that epitope on PstS-1 protein which is anchored on immune cells is accessible to C8 Nb for binding. As compared to rPstS-1 protein treatment, cells treated with protein mixtures derived from Mtb and BCG showed less tracer binding, which might have caused by less availability of PstS-1 protein in protein mixture as well as competition of other ligands from the bacterial lysates for PRR binding. Tracer binding in cells treated with Mtb protein mixture shows that the PstS-1 protein was able to bind macrophage surface receptors even in the presence of other proteins from protein mixture, which also suggests that PstS-1 protein dissociated from Mtb bacilli can bind to immune cells in the TB granuloma and may serve as binding target for C8 Nb. Cells treated with protein mixtures from *E. coli* and *M. smegmatis* showed low tracer binding which was comparable to BSA treated cells, which shows that the C8 Nb will be able to differentiate TB granuloma from other infections. Further, significant reduction in tracer binding on cells treated with similar antigens and incubated with tracer spiked with unlabelled C8 Nb confirmed the specificity of tracer binding.

Thereafter, we sought to determine which PRR binds PstS-1 protein in such way that the antigen epitope is accessible for C8 Nb binding. Contribution of MR in the binding with PstS-1 protein-C8 Nb complex was assessed by incubating WT THP-1 cells with mannose mixed Mtb protein mixture and tracer [27]. However, presence of mannose in the Mtb protein mixture did not produce any significant difference on the tracer binding, which suggests that either the epitope on MR bound PstS-1 protein is not accessible for C8 Nb binding or MR is sufficiently competed out by other mannosylated molecules from Mtb lysate [43]. Similarly, comparable binding of tracer with rPstS-1 protein treated WT and TLR4 KO THP-1 cells confirms that the epitope on PstS-1 protein bound to TLR4 is not accessible for binding. However, in the case of TLR2 KO THP-1 cells, the tracer binding was reduced to the level of BSA treated cells, which suggest that the PstS-1 protein held by only TLR2 can be recognized by C8 Nb.

TB granuloma, made up of Mtb bacilli with surrounding host immune cells, carrying sufficiently accumulated target antigen will be an ideal model for *in vivo* localization study of nanobody. Additionally, in the TB granuloma, Mtb bacilli released from necrotic macrophage may carry more copies of PstS-1 protein produced to survive in the starving phagosomal niche [28, 44]. Also, in all *in vitro* binding assays performed in this study, we have observed stronger immunoreactivity of C8 Nb with Mtb origin PstS-1 protein, either associated with cell wall or present in protein mixture, as compared to the protein homologue produced by BCG. This binding difference of C8 Nb may have arisen due to the differences in PstS-1 protein expression level or binding affinity. However, as Mtb infected animals cannot be handled outside of BSL-III facility, for *in vivo* study we have used mouse model carrying intramuscularaly injected BCG cells, which represents TB granuloma partially without the layer of host immune cells. During *in vivo* localization study, we found less tracer accumulation in the forearm carrying *M. smegmatis* cells than the other arm inoculated with BCG cells. The tracer retention in the hindlimbs without any bacterial inoculations was lowest than the forearms carrying any bacterial pellet, which shows specificity of localization of C8 Nb.

Though, the initial studies showed tracer localization in Mtb nodule even in the presence of immune response against injected antigens [15–17], in this study, presence of unlabelled C8 Nb, ∼1000 fold excess, in the tracer reduced the binding significantly. Therefore, it will be crucial to study the effect of presence of humoral immune response on *in vivo* tracer accumulation in TB granuloma, as the immunodominant PstS-1 protein is known to induce strong antibody response in TB patients [45]. Also, Malpani and Lee et al. used sonicated Mtb lysate as antigen target, which suggest that antibody based imaging won’t be able to discriminate between live and dead Mtb bacilli [15, 17]. Therefore, to understand the utility of nanobody based molecular imaging in treatment response monitoring and predicting disease relapse, it will be important to evaluate the effect of initiation of drug therapy on the extent of tracer localization in Mtb granuloma.

## Materials and Methods

### Materials and reagents

All enzymes including EcoRI, HindIII and T4 DNA Ligase were purchased from New England Biolabs. For nanobody display library construction, T7Select® 1-1 cloning Kit (Cat. No. 70010) was purchased from Merck Millipore. Mouse anti-T7 tail fiber mAb (anti-T7TF, Cat. No. 71530) was procured from Merck Millipore. Rabbit anti-camelid V_H_H antibody-HRP conjugate (Cat. No. A01861) and rabbit anti-camelid V_H_H cocktail-iFluor555 (Cat. No. A02020) were purchased from Genscript.

### Antigen preparation

Entire secreted proteins of laboratory strain Mtb H37Rv were prepared by concentrating the culture filtrate and used for camel immunization. Briefly, Mtb was inoculated in Middlebrook 7H9 broth enriched with ADC growth supplement and 0.05% Tween 80 and grown at 37°C as shaking culture. Exponentially growing Mtb biomass was harvested and washed with phosphate buffered saline (1x PBS). Subsequently, Mtb was inoculated at the density of 10^6^ bacilli/ml in synthetic Sauton’s medium enriched with 0.5% glucose, 0.5% sodium pyruvate, and cultured on an orbital shaker at 37°C [46]. After 14 days of growth, culture was centrifuged at 12,800 × g for 30 min at 4°C. The clear supernatant was collected and saturated with ammonium sulphate (80%). Following overnight incubation at 4°C, protein precipitate was spun down and resuspended in 50ml 1x PBS. Further, insoluble debris were removed by centrifugation and the clear supernatant was filter sterilized through 0.22µm membrane. Protein preparation was concentrated as well as 1x PBS washed on 10kDa cutoff membrane concentrator [47–49]. To ascertain sterility, 100µl protein sample was streaked on Middlebrook 7H10 media and plates were incubated at 37°C for 3 weeks.

### Camel immunization and nanobody library construction

T7 phage displayed immune nanobody library against secretory proteins of Mtb was constructed using the methodologies as reported previously [50]. Briefly, a healthy dromedary maintained at NRCC, Bikaner, India was immunized with Mtb proteins emulsified in Freund’s complete adjuvant (500μg/dose), followed by four boosters prepared in Freund’s incomplete adjuvant at an interval of 21 days. On the seventh day post last booster, camel blood was drawn from jugular vein and PBMC were isolated using Histopaque-1077 according to the manufacturer’s instructions. All the procedures where animal handling was involved, were performed by professional veterinarians as per the guidelines of Institutional Animal Ethics Committee (sanction number: NRCC/PME/6(141)/2000-Tech/Vol-IIPBMC). Camel PBMC pellet was dissolved in 5ml TRIzol reagent and stored at -80°C until further use. Presence of heavy chain only antibodies specific to PstS-1 protein in the camel serum was analysed through ELISA. Total RNA was extracted from PBMCs and subjected to cDNA synthesis by reverse transcription using Hyperscript First strand synthesis kit. Camel produces both conventional (IgG1) as well as heavy chain only antibodies (IgG2 & IgG3), therefore, two step PCR approach was adopted to amplify and separate V_H_H gene fragments from VH amplicons. In the first step, IgG-PCR was performed on cDNA using published primers, i.e., CH2FORTA4 and VHBACKA6, which anneals to the FR1 region and the CH2 domains of all IgG subclasses. After IgG-PCR, amplicons of all IgG subclasses were separated by agarose gel electrophoresis. Due to presence of CH1 domain, the IgG1 subclass produces 900bp size amplicons, while IgG2 & IgG3 subclasses lacking CH1 domain produces amplicons of 690bp & 620bp, respectively, where the length difference arises mainly due to variation in the length of hinge coding region. In the second step, amplicons of IgG2 and IgG3 were used as DNA template in V_H_H PCR to amplify the nanobody coding regions between FR1-FR4 (Primers-BACKHIII and FORECORI). V_H_H amplicons were double digested with EcoRI & HindIII restriction endonucleases. Subsequently, 11ng of V_H_H amplicons and 500ng of T7 phage vector DNA arms (3:1 molar ratio) with cohesive ends were incubated overnight with T4 DNA Ligase at 16°C. A PCR based on primers specific to vector arms (Primers: T7SelectUp & T7SelectDown) was performed to confirm the ligation. For *in vitro* phage packaging, 5μl of ligation reaction mixture was mixed with 25μl of T7 phage packaging extract and incubated for 2 h at 22°C. Phage packaging reaction was terminated by adding 270μl of LB medium and number of assembled infective phages was determined by plaque assay. Before bio-panning, to express and display nanobody on T7 phage capsid, whole library was amplified in liquid culture and stored at -80°C.

### Enrichment and selection of nanobodies specific to PstS-1 protein

High affinity nanobodies against PstS-1 protein were enriched and screened by the bio-panning strategy reported earlier from our lab [50]. Nanobody library was enriched and screened against rPstS-1 protein coated in polystyrene microtiter well plate. For bio-panning, 100µl of rPstS-1 protein (2μg/ml in 0.1M bicarbonate buffer, pH 9.6) was incubated in microtiter wells overnight at 4°C. Wells were washed three times with 1x PBS and then blocked with 200μl of 2% BSA solution for 30 min at 37°C. Phage library was mixed with equal volume of 4% BSA for 15 min at room temperature (RT) and 100μl of library was added to wells for 2 h at 37°C. Further, wells were washed 10 times with 1x PBS containing 0.05% Tween 20 (1x PBST) and bound phages were either eluted in 100μl of 1% SDS for enumeration by plaque assay or amplified *in situ* by adding fresh culture of host *E. coli* Rosetta-gami B5615. In each cycle, BSA coated wells were processed similarly as negative control. To increase washing stringency for enrichment of strong binders, 0.5M and 1M NaCl wash were given following 1x PBST washes in 2^nd^ and 3^rd^ cycles of bio-panning, respectively. After 3 rounds of enrichment, monoclonal phage lysates were prepared for randomly selected 30 nanobody clones. Immunoreactivity of nanobodies was assessed using a two-step screening approach. In the first assay, phage lysates were incubated with the coated antigen, while in the second assay, phages captured via their tail using coated anti-T7 tail fiber antibody, were reacted with ^125^I-labelled rPstS-1 protein. For the first assay, 300μL of rPstS-1 protein was coated overnight into polystyrene star tubes at 4°C. Subsequently, excess rPstS-1 protein solution was discarded and tubes were blocked with 2% BSA solution. Further, monoclonal phage lysates were blocked with equal volume of 4% BSA and 300μl of each clone was added to rPstS-1 coated tubes in triplicate for 2 h at 37°C. Tubes were washed thrice with 1x PBST and incubated with 300μl of ^125^I-labelled anti-T7TF mAb (∼1 lakh CPM) at RT. After 2 h of incubation, tubes were washed thrice with 1x PBST and bound tracer was measured in γ-photon counter. Ten selected nanobody clones were subjected to next assay, where 100μl of each phage lysate was incubated in detachable wells coated with anti-T7TF mAb for 2 h at 37°C. Wells were washed thrice with 1x PBST and then reacted with 100μl of ^125^I-labelled rPstS-1 protein (∼1 Lakh CPM) at RT. After 2 h of incubation, wells were washed thrice with 1x PBST and bound activity was measured in γ-photon counter. Subsequently, inserts of nanobody gene were amplified using PCR (Primers: T7SelectUp & T7SelectDown) and sequenced by sanger sequencing. After nucleotide sequence analysis, nanobody clone 8 (C8 Nb) was selected for further studies.

### Cloning, Overexpression and Purification of C8 Nb

C8 Nb was overexpressed in *E. coli* BL21(DE3) and purified using multiple protein purification methods. Codon-optimised 399bp gene encoding C8 Nb with C-terminal 6xHis tag was synthesized and cloned into pET-22b (+) expression vector. The plasmid construct was electroporated into *E. coli* BL21(DE3) and transformants were selected on ampicillin (100µg/mL) added media. Well isolated colony was inoculated in 10ml Terrific broth supplemented with ampicillin and cultured overnight at 37°C. One litre of Terrific broth inoculated with overnight grown 5ml inoculum was cultured at 37°C and once the OD_600_ of culture reached to ∼1, the protein expression was induced overnight with 1mM IPTG at 20°C. Culture was centrifuged at 2000 x g for 15 min at 4°C and bacterial pellet was resuspended in 40ml lysis buffer (5mM imidazole, 50mM phosphate and 300mM NaCl, pH 8) supplemented with 1mM PMSF and lysozyme (0.2mg/ml). Bacterial suspension was probe sonicated (5 sec on/off) on ice for 30 min. Lysate was centrifuged at 20,000 x g for 45 min at 4°C and clear supernatant was loaded on Ni-NTA agarose column equilibrated with lysis buffer. After washing the column with lysis buffer, bound protein was eluted in 300mM imidazole. Selected protein fractions containing C8 Nb were concentrated on 10kDa cut-off concentrator and resuspended in buffer (10mM NaCl, 20mM Tris pH, 7.8) compatible with anion exchange chromatography. Protein sample was loaded on equilibrated DEAE cellulose column and unbound C8 Nb (pI 7.8) from flow through was concentrated. To remove remaining impurities, protein sample was gel filtered through Superdex^TM^ 75 Increase 10/300 GL column equilibrated with 1x PBS pH 7.4 and C8 Nb fractions were stored at 4°C. After each purification step, nanobody purity was checked on 14% SDS-PAGE.

### Determination of equilibrium dissociation constant (K_D_) using saturation binding ELISA

Microtiter wells were incubated with 100μl of rPstS-1 protein (2μg/ml, pH 9.6) for 1 h at 37°C. After three 1x PBS wash, wells were blocked with 2% BSA for 30 min at 37°C. Wells were then washed twice with 1x PBST and incubated with various concentrations (2pM-2μM) of serially diluted C8 Nb (in blocking solution) for 1 h at 37°C. BSA coated wells were processed similarly as negative control. After 4 1x PBST wash, wells were incubated with 1:10,000 diluted rabbit anti-camelid V_H_H antibody-HRP conjugate for 1 h at 37°C. Subsequently, after five 1x PBST wash, 100µl ready to use TMB substrate was added to the wells and plate was incubated at RT in dark. Color development was stopped with 100µl 1N HCL and absorbance was measured at 450nm.

### Binding kinetics of C8 Nb by Surface Plasmon Resonance

Kinetics of binding between C8 Nb and rPstS-1 protein were analysed on Biacore X100 SPR system (GE Healthcare) at 25^°^C. Briefly, rPstS-1 protein was covalently immobilized on CM5 sensor chip using standard amine coupling. C8 Nb diluted in running buffer [10mM HEPES pH 7.0, 100mM NaCl, 0.005% P20] was then injected onto the antigen bound sensor chip at the increasing concentrations (ranging from 5nM to 240nM) in multi-cycle kinetic assays. Sensor chip was regenerated with glycine buffer and binding kinetics were repeated thrice on the same chip. The acquired data were processed using Biacore evaluation software V2.0.2 (GE Healthcare), and the kinetic rate constants (ka and kd) and equilibrium dissociation constant (K_D_) were determined using kinetic analysis utilizing the SPR data obtained for nanobody concentration such as 5nM, 10nM, 20nM, 40nM and 60nM, where the sensorgrams have sufficient curvature.

### Binding specificity evaluation by Immunoblotting

Mycobacterial strains (Mtb H37Rv, BCG (Mosco) and *M. smegmatis)* were cultured in Middlebrook 7H9 broth added with ADC growth supplement and 0.05% Tween 80 at 37°C. *E. coli* was cultured in terrific broth at 37°C in shaker incubator. Bacterial cells were harvested from log phase cultures and two 1x PBS wash were given. Cells resuspended in 1ml 1x PBS were then transferred to lysing matrix B and lysed in FastPrep-24™ 5G bead beating grinder. After centrifuging the lysate at 20,000 x g for 45 min, supernatant was collected and protein concentration was determined by Bradford assay. Protein samples (2μg rPstS-1 protein or 100μg of crude protein mixture from Mtb, BCG, *E. coli* and *M. smegmatis*) were electrophoresed through 14% SDS-PAGE and then electroblotted on PVDF membrane. Blotted membrane was blocked overnight in 5% skimmed milk powder prepared in 1x PBS. Membrane was subsequently incubated with C8 Nb (2μg/ml in 2% BSA) at RT. After 3 h of binding, membrane was washed (3 x 5ml 1x PBST for 15 min) and incubated with rabbit anti-camelid V_H_H antibody-HRP conjugate (1:10,000) for 2 h at RT. Further, membrane was washed (3 x 5ml 1x PBST for 15 min) and developed with chemiluminescent substrate.

### ^125^I labelling of C8 Nb

Radioiodination of C8 Nb was done using Iodogen method as described previously [51]. To prepare Iodogen coated tubes, 100µl of Iodogen (1mg/ml in CHCl_3_) was added to 1.5ml Eppendorf tubes and air dried to form Iodogen layer on tube wall. Reducing agent free iodination grade Na^125^I was supplied by BRIT, Mumbai, India. Fifty microliters of C8 Nb (100µg/ml in 0.5M phosphate buffer, pH 7.4) was dispensed to the bottom of Iodogen coated tube. Further, 4µl of ^125^I (∼771µCi) activity was added to the same tube and after gentle mixing, the tube was incubated for 5 min at RT. Labelling was terminated by adding 200µl of 50mM phosphate buffer and reaction mixture was loaded on Sephadex G-25 coarse column. Subsequently, 2% BSA solution in 1x PBS was passed through the column, and elution fractions of 0.5mL were collected.

### ^125^I-C8 Nb Radioimmunoassay

100μl of rPstS-1 protein (2μg/ml, pH 9.6) was coated overnight in detachable microtiter wells at 4°C. After 1x PBS wash, wells were blocked with 200μl of 2% BSA for 30 min at 37°C. Wells were subsequently incubated with 100μl of tracer (^125^I-C8 Nb, ∼119000 CPM). Tracer was used either in its normal form or spiked with unlabelled C8 Nb (15μg/ml). BSA coated wells were treated similarly as negative control. After 1 h incubation at RT, wells were 4 times washed with 1x PBST and bound tracer was measured in γ-photon counter.

### Immunoreactivity of ^125^I-C8 Nb against PstS-1 protein present in the cell wall of drug sensitive as well as MDR mycobacteria

Mycobacteria such as Mtb H37Rv, Mtb LAM, Mtb Beijing, BCG (Mosco) and *M. smegmatis* were cultured in Middlebrook 7H9 broth enriched with ADC growth supplement and 0.05% Tween 80 at 37°C in shaking culture. *E. coli* was cultured in terrific broth at 37°C in shaking condition. Log phase cultures of mycobacteria were harvested and passed through 26 gauze needle. Two 1x PBS wash were given to single cell suspensions and OD_600_ was adjusted to 1. One millilitre of each strain aliquoted in 1.5ml Eppendorf tube was centrifuged and pellet was resuspended in 0.5ml 2% BSA at 37°C. After 30 min blocking, cells were spun down and resuspended in 100μl of ^125^I-C8 Nb (∼1,20,000 CPM) prepared in 2% BSA. Similar bacterial pellets were also treated with tracer spiked with unlabelled C8 Nb. After 30 min binding at RT, cells were pelleted down and washed twice with 1x PBS. Further, 0.5ml 10% formaldehyde was added on the top of pellets and bound tracer was measured in γ-photon counter.

### Immunoreactivity of ^125^I-C8 Nb with rPstS-1 protein adhered on macrophages

RAW 264.7 cell line was cultured under 5% CO_2_ in RPMI 1640 medium supplemented with 10% FBS. Confluent cells were trypsinzed and 0.25×10^6^ cells resuspended in 1ml complete medium were seeded in 12 well cell culture plate. Next day, cells were washed with 1x PBS and fixed in 4% formaldehyde solution for 15 min at RT. Then, wells were thrice washed with 1x PBS and blocked with 2% BSA for 30 min at 37°C. Further, cells were incubated with protein samples (200µg of crude protein mixture prepared from bacterial lysates of Mtb, BCG, *M. smegmatis*, *E. coli* or 2µg rPstS-1 protein) resuspended in 200µl of blocking solution for 1 h at 37°C. Wells were washed thrice with 1x PBS and 200µl of ^125^I-C8 Nb (∼2×10^5^ CPM) was added in each well. After incubating for 1 h at 37°C, cells were washed thrice with 0.5ml 1x PBS and 200µl of 1% SDS was incubated in each well for 5 min at RT. Well contents were collected in tubes and bound tracer was measured in γ-photon counter.

### Determination of PRR involved in macrophage binding with PstS-1 rotein-C8 Nb complex

Wild type THP-1 cells were cultured under 5% CO_2_ in RPMI 1640 medium supplemented with 10% FBS at 37°C. TLR-2 and TLR-4 KO THP-1 cell lines were cultured and maintained in RPMI 1640 medium under selection pressure of Blasticidin (10µg/ml) and Zeocin (100µg/ml). Approximately 0.5×10^6^ cells resuspended in 1ml complete medium containing phorbol myristate acetate (PMA) (100ng/ml) were seeded in 12 well cell culture plate and incubated at 37°C for overnight. Next day, cells were washed twice with 1x PBS and fixed with 4% formaldehyde for 15 min at RT. In the next step, cells were washed thrice with 1x PBS and incubated with 0.5ml 2% BSA for 30 min at 37°C. Further, blocking solution was discarded and cells were incubated with 2µg of rPstS-1 protein resuspended in 200µl of blocking solution for 1 h at 37°C. Subsequently, wells were washed thrice with 1x PBS and 200µl of ^125^I-C8 Nb (∼2.3×10^5^ CPM) was incubated in each well. Similarly, to evaluate the role of MR in binding, WT THP-1 cells were treated with 200µg of Mtb protein mixture (with or without mannose, 1mg/ml), resuspended in 200µl 2% BSA. Spiked tracer was prepared by mixing unlabelled C8 Nb (15µg/ml) and added to similarly treated control wells. After 1 h binding at 37°C, cells were washed thrice with 1x PBS and 200µl of 1% SDS was added in each well for 5 min at RT. Further, well contents were collected and bound activity was measured in γ-photon counter. Similarly, for fluorescence microscopy, cells were cultured and prepared in µ-slide 8 well glass bottom plate. WT as well as KO THP-1 cells treated with rPstS-1 protein (10µg/ml) were incubated with 100µl of C8 Nb (1µg/ml). After 1 h binding at 37°C, cells were washed thrice with 1x PBS and incubated with 100µl of 1:500 diluted rabbit anti-camelid V_H_H cocktail-iFluor555 for 1 h at 37°C. Further, cells were washed thrice with 1x PBS and incubated with 100µl of DAPI stain (1µg/ml in 1x PBS) for 10 min at RT. Subsequently, after 1x PBS wash, 100µl 2% formaldehyde was added to wells and images were acquired on Leica Mica Microhub.

### In vivo localization of ^125^I-C8 Nb around BCG cells

For *in vivo* localization study, 4-5 week old BALB/c mice were obtained from animal house, BARC, Trombay and maintained at animal house, RMC, Parel. Three days prior to the study, drinking water was replaced with potassium iodide solution (1g/L water). Log phase cultures of *M. smegmatis* and BCG were harvested and passed through 26 gauge needle and kept standing for 10 min to remove bacterial clumps. Bacterial suspensions were washed twice with 1x PBS and 5×10^8^ cell number was adjusted in 50µl of 1x PBS. Animals were anesthetized with ketamine (IP) and 50µl of BCG and *M. smegmatis* cell suspensions were injected through intramuscular route in right forelimb and left forelimb, respectively. Further, 75µl of ^125^I-C8 Nb (∼150ng C8 Nb, 25.91µCi) was injected through catheterized tail vein. After 4 h, animals were euthanized with overdose of anaesthesia and limbs were collected for γ-photon counting.

## Acknowledgements

We thank Bhabha Atomic Research Centre, DAE, Govt. of India, for funding this project, and National Research Centre for Camel – ICAR for collaboration and providing necessary support in camel maintenance, immunization and booster. We thank Dr. Yogita Shete, Radiation Medicine Centre; for help in conducting animal experiments. We thank Mr. U. F. Shaikh for assistance in biodistribution study.

## Author contributions

YPD, KJ, SK and PKG conceived the study and designed all experiments and analysed the data. All cloning, protein purification and in vitro assays were performed by YPD. Camel immunization and blood collections were carried out by SKG and KJ. SPR data collection and analysis were done by GDG and SS. Manuscript draft prepared by YPD was reviewed and edited by KJ, SK and PKG.

## Abbreviations

Nb: Nanobody
V_H_H: Variable domain of the heavy chain of the heavy-chain only antibody
VH: Variable domain of the heavy chain of the conventional antibody
IgG: Immunoglobulin G
SDS-PAGE: Sodium dodecyl sulfate polyacrylamide gel electrophoresis
Fabs: Antigen-binding fragments
PBMCs: Peripheral blood mononuclear cells
PFU: Plaque-forming units
BSA: Bovine serum albumin
E. coli: Escherichia coli
HRP: Horseradish peroxidase
ELISA: Enzyme-linked immunosorbent assay
SPR: Surface plasmon resonance
IPTG: Isopropyl-β-d-thiogalactoside
6xHis: Hexahistidine
pH: Potential of hydrogen
mAbs: Monoclonal antibodies
FBS: Fetal bovine serum
K_D_: Equilibrium dissociation constant
Ka: Association rate constant
Kd: Dissociation rate constant
OD_600_: Optical density at 600nm wavelength
ADC: Albumin dextrose catalase
CPM: Counts per minute

